# Regional connectivity and partial residency of capelin (*Mallotus villosus*) in the Gulf of St. Lawrence inferred from otolith chemistry

**DOI:** 10.64898/2026.06.18.733096

**Authors:** Romaric Jac, Elisabeth Van Beveren, Olivier Le Pape, Mathieu Boudreau, Lola Coussau, Pascal Sirois, Dominique Robert, Pablo Brosset

## Abstract

Capelin (*Mallotus villosus*), a key forage fish in the Northwest Atlantic, links zooplankton to predators including commercial fishes, seabirds, and marine mammals, yet its life-cycle movements in the Gulf of St. Lawrence (GSL) remain poorly understood. Between 2022 and 2024, otoliths from 927 individuals collected during and after spawning were analysed by Laser Ablation Inductively Coupled Plasma Mass Spectrometry (LA-ICP-MS). Building on previous work on regional structuring, seven trace elements (Li, B, Mg, K, Zn, Sr, Ba) were used to discriminate three regions. Early-life regional signatures were inferred through an edge-to-core approach, assigning otolith core chemistry to one of these regions using quadratic discriminant analysis. The core was treated as an integrated early-life signal (late-larval to early juvenile period) rather than a strictly natal signature. Spatial variation in core chemistry was consistent across cohorts, mirroring the stability documented on the otolith edge. Results revealed widespread dispersal alongside partial regional residency: individuals with northeastern early-life signatures showed the strongest correspondence between early-life and capture regions, whereas other regions were more connected. Fish sampled during spawning were more often reassigned to their inferred early-life region than post-spawning fish, a regional-scale homing-like pattern consistent with regional spawning fidelity. This coexistence of dispersive and resident strategies likely generates a portfolio effect buffering the population against environmental variability and localised reproductive failures.

## Introduction

Forage fishes play a pivotal role in marine ecosystems worldwide, linking plankton production to higher trophic-level predators such as seabirds, marine mammals, and piscivorous fishes (Cury *et al*., 2011; Ruzicka *et al*., 2013). Characterised by short lifespans, rapid growth, and strong dependence on secondary production, these small pelagic species often undergo pronounced “*boom-and-bust*’’ dynamics, with periods of high abundance followed by abrupt declines (Baumgartner *et al*., 1992; McClatchie *et al*., 2017). These natural fluctuations linked to environmental variability can be amplified by anthropogenic pressures, occasionally leading to stock collapses or long-term reductions in productivity (Peck *et al*., 2021; Szuwalski & Hilborn, 2015; Szuwalski *et al*., 2023). A key pathway through which these pressures affect population trajectories is the spatial and temporal distribution of individuals, making it essential to understand habitat use and movement ecology across life stages, to reduce uncertainties in stock assessment, inform ecosystem-based strategies, and anticipate climate-driven changes in marine ecosystems (Boldt *et al*., 2022; Peck *et al*., 2021). Despite recent advances in monitoring and analytical methods, substantial gaps remain in the capacity to reveal the movements, migrations, and life-history connectivity of forage species (Boldt *et al*., 2019, 2022; Jac *et al*., 2025).

Among forage fish species, capelin (*Mallotus villosus* O.F. Müller, 1776) is one of the most important in Canadian waters (Buren *et al*., 2019), representing the main prey of several commercially important fishes, seabirds and marine mammals (e.g., Brown-Vuillemin *et al*., 2022; Morissette *et al*., 2006; Koen-Alonso *et al*., 2021). In these waters, capelin occur primarily off Newfoundland and Labrador and throughout the Gulf of St. Lawrence (GSL), with smaller populations in the Canadian Arctic and in the Pacific Ocean off British Columbia (McNicholl *et al*., 2016; McQuinn *et al*., 2012; Simard *et al*., 2002). Capelin undertake extensive seasonal migrations, particularly during the spawning period when they move from offshore overwintering areas to inshore spawning habitats (e.g., Nakashima & Wheeler, 2002; Templeman, 1948). Spawning generally begins in early summer, although sea temperature, ice cover, and other local conditions generate substantial inter-annual and regional variability (Buren *et al*., 2014; Carscadden *et al*., 1997; Penton *et al*., 2012). Although weak genetic structuring and evidence of local adaptation have been reported between the GSL and the Newfoundland-Labrador Shelf (Cayuela *et al*., 2020; Kenchington *et al*., 2015), gene flow appears sufficient for these regions to be managed as a single unit (Boudreau *et al*., In press).

In contrast to regions where substantial research effort has been devoted to capelin, particularly off Newfoundland, and Labrador (e.g., Davoren *et al*., 2015; Maxner *et al*., 2016; Murphy *et al*., 2021; Tripp & Davoren, 2025; Tripp *et al*., 2020, 2024), the GSL lacks both dedicated scientific surveys and region-specific studies. In this area, the spawning season is spatially structured, beginning in the upper estuary and progressing eastward and northward through July and August (Carscadden *et al*., 1997), and capelin undertake spawning migrations from deeper waters to nearshore habitats (Boudreau *et al*., In press). However, limited knowledge of migration routes, regional connectivity, and spatial structuring continues to hinder accurate estimation of spawning-stock biomass and to constrain robust stock assessment and management (Boudreau *et al*., In press; Jac *et al*., 2025).

A wide range of approaches, including tagging, stable isotope and trace-element analyses, genetic markers and environmental DNA can provide insights into stock structure and movements (Jac *et al*., 2025; Randon *et al*., 2021; Bergman *et al*., 2026). Among available tools, otolith chemistry has proven particularly effective for discriminating capelin from different regions and reconstructing their movements (Davoren *et al*., 2015; Fink-Jensen *et al*., 2022; Lazartigues *et al*., 2016; Maxner *et al*., 2016; Tripp *et al*., 2020). Otoliths are metabolically inert, incrementally growing structures that incorporate trace elements from surrounding waters, creating a chronological archive of environmental exposure (Campana, 1999; Campana & Neilson, 1985). The otolith core records conditions experienced during early life, whereas the edge reflects the environment at capture, allowing reconstruction of individual movements and inference of population connectivity (Campana, 1999; Elsdon *et al*., 2008; Sturrock *et al*., 2015). Using otolith-edge chemistry, Jac *et al*., (2026) identified three hydrographically distinct regions within the GSL, characterised by multi-year stability in chemical fingerprints: the Estuary, the South, and the Strait of Belle Isle (SBI). Building on this work, the present study aims to (i) characterise the early-life regional chemical signature recorded in the otolith core and assign individuals to one of these regions, using an edge-to-core method, (ii) compare inferred early-life regions with capture regions to describe regional connectivity and (iii) investigate the seasonal and biological factors associated with movement between regions.

## Materials and methods

### Study area and capelin collection

The study was conducted in the St. Lawrence Estuary and Gulf (hereafter GSL; Fig 1), a semi-enclosed system characterised by complex bathymetry and oceanography (Galbraith *et al*., 2025). The southern GSL consists of a shallow plateau (∼80 m), whereas the northern GSL is dominated by deep channels and continental slopes (mean depth ∼240 m; max ∼500 m). In summer, the water column exhibits a three-layer thermal structure comprising a warm surface layer, a cold intermediate layer (CIL; 50-120 m), and a deeper Atlantic-origin layer with intermediate temperature (Galbraith *et al*., 2025). The CIL represents a key thermal habitat influencing capelin distribution (Chamberland *et al*., 2022).

**Fig 1.**
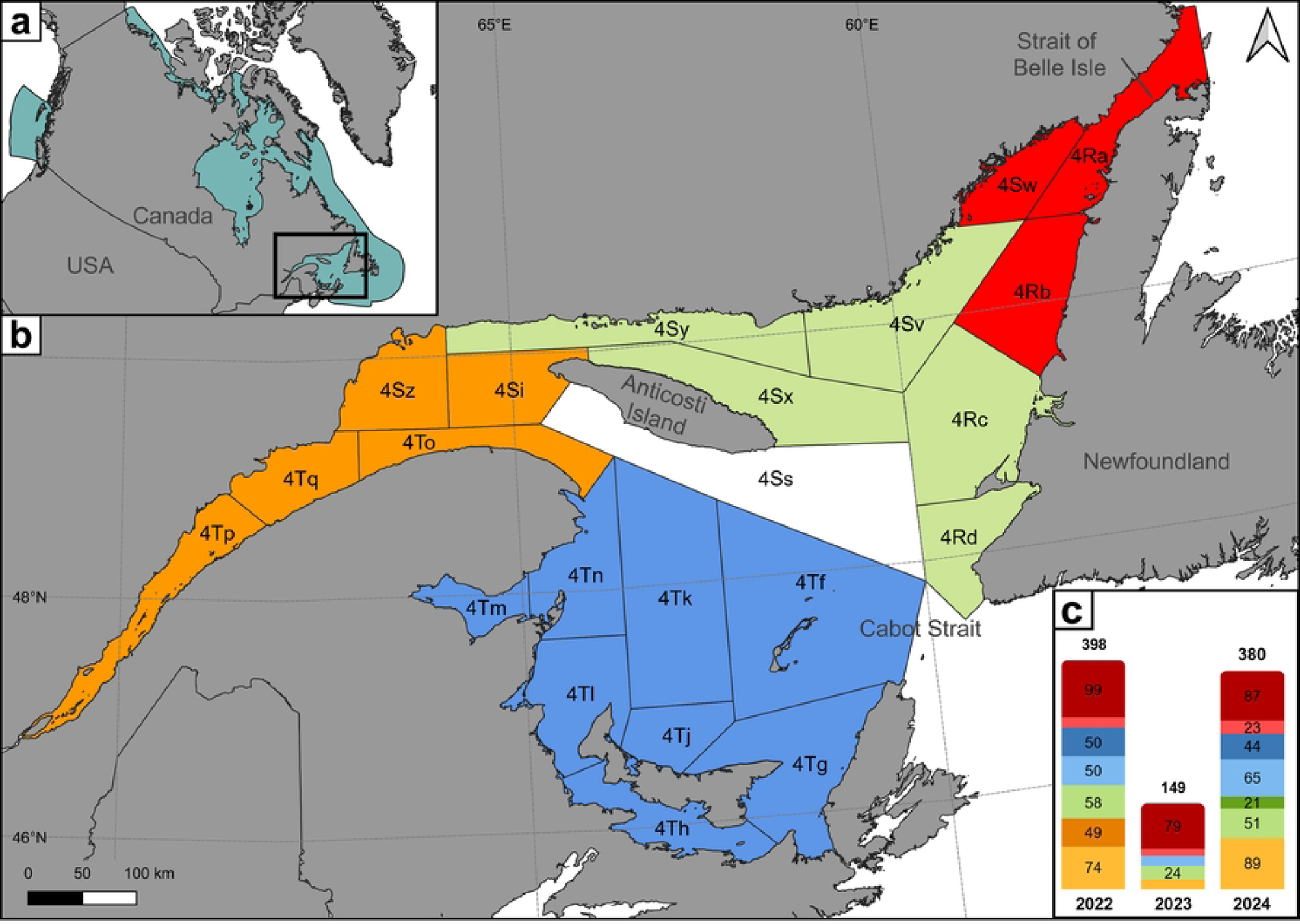
Distribution and sampling of capelin across the study area. (a) Schematic distribution of capelin across Canada’s three oceans, based on Jac *et al*. (*2025*) and (b) the study area with the NAFO sub-divisions and subdivided into four regions after the clustering analysis; (c) a tacked bar chart displaying the number of individuals sampled per year and region. Colours represent regions as follows: orange for the Estuary, green for the North, blue for the South, and red for the Strait of Belle Isle. In the stacked bar chart, light shades represent samples from DFO multi-species bottom trawl surveys, while dark shades correspond to commercial fishery samples. Sample sizes are labelled when exceeding 20 individuals. Shapefiles used for map construction were obtained from Runfola et al. (2020). Map projections: (a) ESRI:102001, Canada Albers Equal Area Conic; (b) EPSG:32198, NAD83 / Québec Lambert.

From 2022 to 2024, 927 capelin – 398 in 2022, 149 in 2023, and 380 in 2024 – were collected across the GSL. Individuals were selected using a stratified random design to minimise potential biases, balancing sampling period (during/after spawning), Northwest Atlantic Fisheries Organization (NAFO) subdivision (Fig 1), fish length, and sex. Fork length (FL) was measured, and sex was determined by external examination or gonad inspection when necessary. The dataset comprised 505 females, 356 males, and 66 individuals of undetermined sex, ranging from 7.5 to 19.0 cm FL.

Fish originated from two sources: commercial fisheries and scientific surveys (Fig 1). Commercial fishery samples were collected between May and July through Fisheries and Oceans Canada (DFO) port sampling, coinciding with the spawning period (Boudreau *et al*., In press). These samples were obtained from nearshore fisheries using pound nets, traps, and purse seines, with spatial resolution limited to Northwest Atlantic Fisheries Organisation (NAFO) subdivisions, primarily 4Ra, 4Sw, and 4Tn (Fig 1). Commercial samples mainly comprised larger individuals (10.5-19 cm FL). Scientific samples were obtained from DFO’s annual multispecies bottom trawl surveys conducted in August-September, corresponding to the post-spawning period (Chamberland *et al*., 2022). These surveys used a standardised otter trawl and provided precise spatial coverage across the GSL. Although sampling methods differed between sources, these differences do not affect the study objectives, as analyses focus exclusively on otolith elemental chemistry, which reflects environmental exposure rather than capture conditions (Jac *et al*., 2026).

### LA-ICP-MS otolith chemistry and data processing

Detailed otolith preparation and LA-ICP-MS procedures are described in Jac *et al*., (2026). Briefly, sagittal otoliths were extracted following standard protocols (Campana, 1999), and only the right otolith of each individual was analysed. Otoliths were rinsed with ultrapure water, gently cleaned, air-dried, and mounted on microscope slides using thermoplastic glue. They were ground and polished using successive abrasive papers (1200 µm, 5 µm, and 1 µm) with circular and figure-eight motions. This two-dimensional polishing approach reliably exposes the ventral edge of capelin otoliths and has been validated in previous studies (Lazartigues *et al*., 2014, 2016). Elemental analyses were conducted at LabMaTer (Université du Québec à Chicoutimi, Canada) using Laser Ablation Inductively Coupled Plasma Mass Spectrometry (LA-ICP-MS; Resolution M-50 Excimer 193 nm laser coupled to an Agilent 7900x qICP-MS). For each otolith, a single continuous laser transect was ablated along the ventral growth axis, extending from the core to the outer margin (Fig 2). This approach captures the full chronological sequence of elemental incorporation across the otolith, from early life to capture. Ablation parameters were: fluence 5 J·cm⁻², repetition rate 25 Hz, beam diameter 44 µm, and scan speed 5 µm·s⁻¹. This beam size optimised signal stability and material removal while maintaining adequate temporal resolution for juvenile and adult capelin (Lazartigues *et al*., 2014). Data were acquired at 4 measurements·s⁻¹ (integration time 0.252 s). Ablated material was transported using an argon-helium gas mixture with added nitrogen, and each transect was preceded by a 30-s gas blank. A total of 38 isotopes were measured. Certified reference materials (NIST SRM-610/612; USGS GSE-1, GP4-A, MACS-3) were analysed at the beginning and end of each session and after every seven otoliths to ensure analytical accuracy and precision. Measured element-to-calcium ratios were within acceptable limits relative to published reference values (USGS; Weis *et al*., 2022). Data reduction was performed using Iolite (Paton *et al*., 2011) in Igor Pro. Calcium (^44^Ca; 38.02% of otolith mass; Campana, 1999) was used as an internal standard, with calibration based on NIST SRM-610. Only otoliths exhibiting stable calcium signals were retained. Element concentrations were expressed in parts per million (ppm), and limits of detection (LOD) were calculated as three times the standard deviation of the gas blank divided by analyte sensitivity (Lazartigues *et al*., 2014).

**Fig 2.**
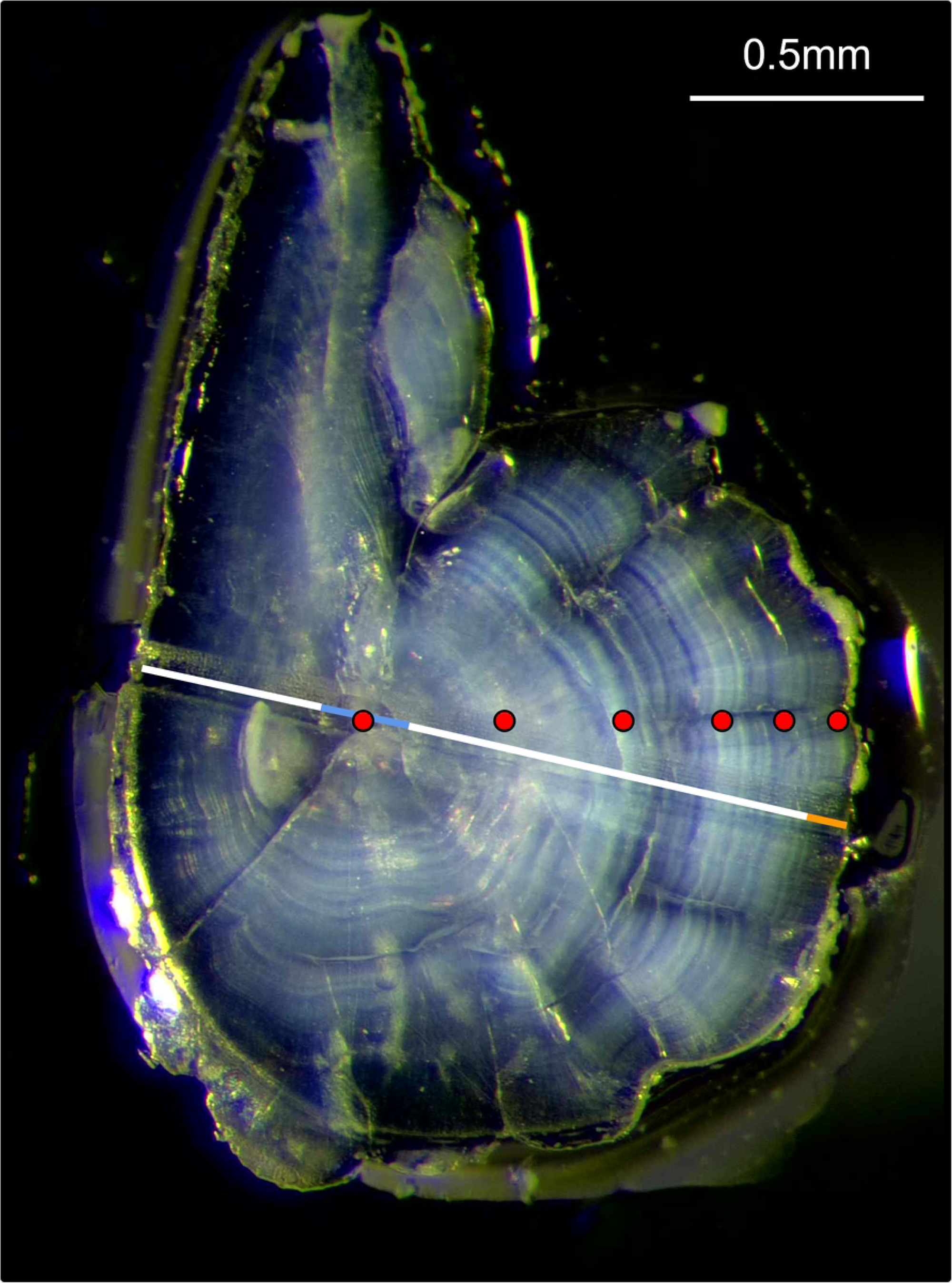
Sanded capelin otolith submerged in water following LA-ICP-MS analysis. The long white line represents the ablation transect performed during the procedure, with the white segment indicating the portion extending from the otolith core to the outer margin. The two short segments mark the core (blue) and the 40μm edge region (orange) analysed in this study. Points denote the centres of winter rings used to determine the individual’s age.

Specific portions of each continuous transect were then extracted to characterise life-stage specific environmental signatures. The core was identified visually and measurements within 20 µm of the otolith core (Fig 2) were used to define an early-life chemical signature, following established approaches for capelin and other forage fishes (e.g., Couillard *et al*., 2022; Lazartigues *et al*., 2016; Coussau *et al*., 2026). Precise targeting of the core remains challenging given the convex otolith morphology, and the beam diameter inevitably integrates surrounding early post-hatch material (Fink-Jensen et al., 2021, 2022). Importantly, the elevated trace-element concentrations that characterise the pre-hatch and post-hatch regions of larval capelin otoliths in the GSL (Coussau *et al*., 2026) and elsewhere were not recovered in the core region of the adult otoliths analysed here. The core is therefore not treated as a strictly natal (hatching) signal but rather as an integrated early-life signature, most plausibly reflecting a late-larval to early-juvenile period. In contrast, the capture signature was defined using the outermost 40 µm of the otolith (Fig 2), which reflects the environment experienced immediately prior to capture (Davoren & Halden, 2014; Fink-Jensen *et al*., 2021; Jac *et al*. 2026).

Seven elements were retained based on their high discriminatory power: lithium (Li), boron (B), magnesium (Mg), potassium (K), zinc (Zn), strontium (Sr), and barium (Ba). The selection of these elements was detailed in Jac et al., (2026) and included validation that spatial patterns were consistent across years. Demonstrating spatial persistence through time is a prerequisite for subsequent analyses, especially given that temporal variability can be considerable and for some elements may exceed variability in ambient water chemistry (Hüssy et al., 2020). This is in part because both non-essential and essential elements were included. Essential elements (e.g., Mg and K) may be influenced by physiological regulation rather than solely reflecting ambient environmental conditions. However, they can still contribute to regional discrimination if such processes exhibit persistent spatial structure. Because inter-annual variation in these physiological processes can be substantial, the temporal stability of these elements as regional discriminators across years may be reduced. This multi-element approach is consistent with previous otolith chemistry studies on capelin, which have similarly incorporated physiologically regulated elements such as Mg, K, and Zn in multivariate analyses (e.g., Tripp et al., 2020; Lazartigues et al., 2016). Accordingly, the selected element suite should be interpreted as markers of regional origin rather than strictly as proxies of environmental conditions. Values below the LOD were replaced with LOD/2, and outliers exceeding ±5 standard deviations were replaced by the mean of adjacent measurements. Elemental concentrations were converted to element:Ca ratios, and non-normally distributed elements (Li, B, Mg, Zn) were log(x+1)-transformed prior to statistical analyses.

### Age readings and age assignment

Capelin age was estimated by counting growth annuli on the same sagittal otoliths used for LA-ICP-MS elemental analysis. Otoliths were photographed using a Leica M125C microscope, and growth increments were measured in ImageJ (Schneider *et al*., 2012; Fig 2). This study represents the second attempt at capelin age estimation in the GSL, following Bailey *et al*., (1977a). Unlike other regions (e.g., Davoren & Halden, 2014; Fink-Jensen *et al*., 2022), interpreting annuli from GSL capelin was challenging due to otolith opacity and the inconsistent presence of the metamorphic check. Consequently, only 307 out of 927 otoliths with clearly visible and interpretable annuli were retained for direct age determination. Age readings were independently performed by two trained readers, with disagreements resolved jointly. Otoliths for which age consensus could not be reached were excluded (n = 5; 1.63%), yielding a final age dataset of 302 individuals (*dataset_age*; Table 1). Ages ranged from 1 to 6 years (Fig 3; S1 Fig), and individuals were distributed throughout the entire study area (S2 Fig).

**Table 1.** Number of individuals per cohort, including fish aged directly from otoliths (*Dataset_age*, n = 302) or supplemented with individuals whose ages were estimated using a probabilistic Age–Length Key (pALK) (*Dataset_age_supplemented*, n = 594).

|  | 2016 | 2017 | 2018 | 2019 | 2020 | 2021 | 2022 | 2023 | Total |
| --- | --- | --- | --- | --- | --- | --- | --- | --- | --- |
| *Dataset_age* | 6 | 23 | 59 | 42 | 79 | 71 | 16 | 6 | **302** |
| *Dataset_age_supplemented* | 9 | 34 | 128 | 130 | 132 | 109 | 46 | 6 | **594** |

**Fig 3.**
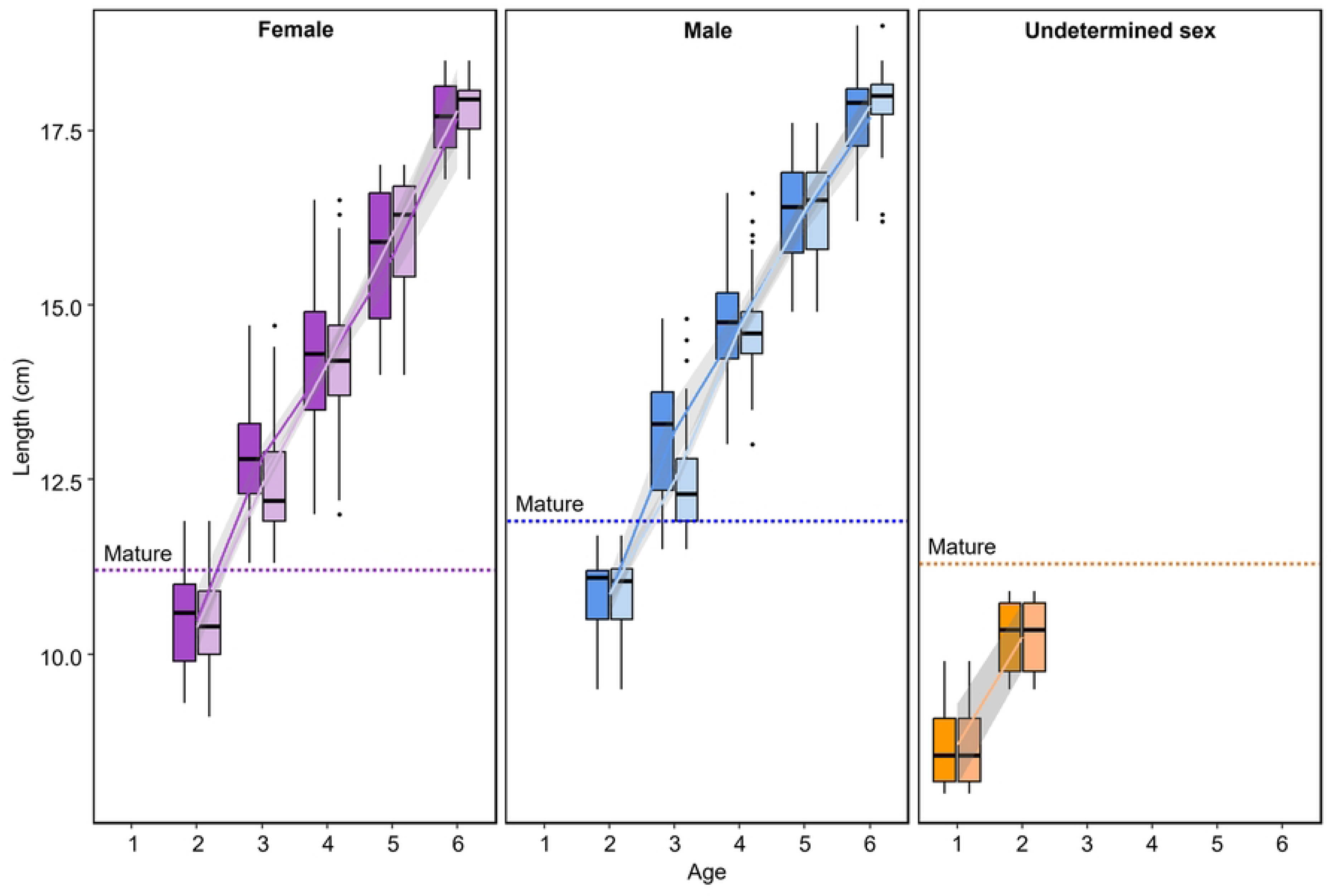
Age-Length relationship for females (pink), males (blue), and individuals of undetermined sex (orange) as well as the age–length profiles 95% confidence intervals. Dark colours indicate individuals aged directly from otoliths (Dataset_age, n = 302), while light colours represent individuals whose ages were estimated using a probabilistic Age–Length Key (pALK) based on multinomial regression (Dataset_age_supplemented, n = 594). An additional 171 females and 121 males were included through the pALK.

To assign ages to unaged individuals, a probabilistic Age-Length Key (ALK) based on multinomial regression (Campana, 2001; Kimura, 1977) was constructed using the 302 directly aged fish. Probabilities of assignment to each age class were estimated using the *multinom* function from the *nnet* package (Venables & Ripley, 2002), generating sex-specific ALKs to account for differences in growth patterns. Unaged individuals were assigned to an age class when the posterior probability exceeded 75%. This threshold yielded an 85.5% success rate in the reassignment of aged individuals, supporting the ALK as a robust tool for assigning ages in population-level analyses. Based on the ageing of an additional 292 fish, the dataset was expanded to 594 individuals (*dataset_age_supplemented*; Table 1; S1 Fig), distributed throughout the entire study area (S2 Fig). Comparison of age-length relationships indicated strong agreement between the observed and ALK-augmented datasets (Fig 3), with the age distributions of newly assigned individuals closely matching those of directly aged fish, showing only minor deviations around age 3.

### Determining early-life region using edge-to-core reassignment

Each individual’s early-life region was inferred using an edge-to-core reassignment method (Avigliano, 2022; Reis-Santos *et al*., 2023), in accordance with the recommendations of Arai *et al*., (2026). Otolith core chemical signatures, defined here as the innermost 20 μm of the transect and representing an integrated early-life environmental signal (Couillard *et al*., 2022; Lazartigues *et al*., 2016; Coussau *et al*., 2026), were assigned to one of three regions using a single multi-class quadratic discriminant analysis (QDA) trained on otolith edge signatures. Otolith edge chemistry represents the most recently deposited material prior to capture and may reflect either spawning habitat or post-spawning habitat, depending on the sampling period. Edge chemistry defined three hydrographically distinct regions within the GSL (Estuary, South, and SBI), based on multivariate analyses of otolith edge elemental composition conducted by Jac *et al*. (2026).

The fitted QDA model estimated posterior probabilities of membership for each otolith core signature across the three regions. Each individual was assigned to the region associated with the highest posterior probability. Assignments were retained when the maximum posterior probability exceeded 50%; a threshold above the random expectation of ∼33% for three regions, ensuring that retained assignments were supported by more than chance alone. Individuals below this threshold were classified as belonging to an ‘undetermined’ region. More conservative thresholds (70% and 90%) were also evaluated. Although a higher threshold increased the proportion of ‘undetermined’ individuals, it did not affect the overall spatial patterns of early-life region assignment (S3 Fig), indicating that the results are robust to the choice of assignment threshold.

Although some inter-annual variability in elemental composition was observed by Jac *et al*. (2026) on the otolith edge, spatial differences among the three regions consistently exceeded temporal variation, supporting the use of elemental signatures as reference classes across cohorts. Despite this consistent inter-annual signal on the otolith edge (Jac *et al*., 2026), the presence of ‘cohort effects’, i.e., differences among year-classes (Campana, 1999; Elsdon *et al*., 2008; Gillanders, 2002, 2005) remained a potential confounding factor in an inter-annual comparison. To assess the potential influence of these ‘cohort effects’, individuals from both datasets (*dataset_age* and *dataset_age_supplemented*) were grouped by cohort (Table 1), and two classification approaches were compared: (i) cohort-stratified, in which individuals were classified within their respective cohort and a separate QDA was fitted for each, and (ii) pooled, in which all individuals were classified together without considering cohort. Consistency between models was expressed as the percentage of individuals assigned to the same early-life region across both approaches.

Inferred early-life regions of individuals caught during spawning and post-spawning were then mapped based on shapefiles from Runfola et al. (2020), allowing spatial visualisation of regional connectivity within the GSL.

### Drivers of movement between regions

Connectivity among regions of the GSL was further examined in relation to potential explanatory factors, including life stage and seasonal timing. Individuals were classified as either residents or migrants. Residents were defined as individuals whose inferred early-life region matched their capture region, whereas migrants were those whose inferred early-life region differed from their capture region. Movement probability was then analysed using a binomial generalised linear model (GLM), with status (migrant versus resident) as the binary response variable. Predictor variables included maturity, sex, sampling timing of capture (spawning versus post-spawning) and inferred early-life region. Maturity status was determined from sex-specific length thresholds: females >11.2 cm and males >11.9 cm (DFO, *unpublished data*). A forward stepwise model selection procedure based on the Akaike Information Criterion (AIC; Akaike, 1974) was used to identify the most parsimonious GLM.

## Results

### Cohort effects on inferred early-life region assignment

First, the influence of cohort on inferred early-life region assignment was assessed. Reassignment between age-stratified and pooled QDA models was highly consistent across regions (Table 2), indicating that cohort has little influence on assignment. Indeed, for the directly measured ages (*dataset_age*), regional concordance ranged from 74.6% to 87.0%, with an overall agreement of 80.5% between the two modelling approaches. In the dataset enriched with ALK-estimated ages (*dataset_age_supplemented*), concordance was higher across all regions, ranging from 95.7% to 97.6%, yielding an overall consistency of 97.0%. This increase likely reflects the larger sample size in the enriched dataset. Collectively, these results indicate that age class exerts a negligible effect on early-life region assignment at the scale of the entire GSL. This concordance reflects the consistency of assignments across cohorts within the present dataset rather than a formal assessment of the temporal stability of core signatures within regions (see Discussion). Consequently, in the absence of a detectable cohort effect, a pooled QDA model without age-class stratification was applied to all eligible individuals (n = 927) for subsequent analyses.

**Table 2.** Concordance of predicted regions by collection region, based on otolith core chemistry using quadratic discriminant analysis (QDA). Comparisons were made using both an age-class–stratified model and a pooled model to evaluate the effect of cohort on assignment outcomes. Analyses were conducted on two datasets: individuals aged directly from otoliths (*Dataset_age*, n = 302) and individuals aged via a probabilistic Age–Length Key (*Dataset_age_supplemented*, n = 594).

|  | Collection regions (%) |  |  |  | Global Consistency (%) |
| --- | --- | --- | --- | --- | --- |
|  | Estuary | South | North | SBI |  |
| *Dataset_age* | 74.6 | 76.6 | 78.0 | 87.0 | **80.5** |
| *Dataset_age_supplemented* | 96.9 | 97.1 | 95.7 | 97.6 | **97.0** |

### Regional patterns of early-life region assignment

Reassignment of otolith core chemistry to one of the three chemical regions, or to ‘undetermined’ when the 50% posterior probability threshold was not met, revealed clear differences in inferred early-life regions among capture regions (Table 3). The SBI was the most frequently assigned region. Individuals captured in the Estuary were most frequently assigned to the SBI region (38.0%), followed by the South region (24.9%) and the Estuary itself (15.7%). Capelin captured in the South were primarily assigned to their own region (31.0%) and to SBI (26.1%), whereas individuals captured in SBI were predominantly reassigned to their own region (70.4%), highlighting stronger regional residency (here, residency refers to the correspondence between capture region and inferred early-life region). The North region appeared highly connected with SBI, with just under half of its individuals assigned an SBI early-life signature. The proportion of individuals with an ‘undetermined’ early-life region varied among regions. While it was relatively low in SBI (11.1%) and North (18.2%), it was higher in the Estuary (21.4%) and South (23.4%), indicating greater uncertainty in assigning individuals captured in these regions (Table 3).

**Table 3.** Reassignment results of capelin otoliths based on edge chemistry, using quadratic discriminant analysis (QDA). Values represent the percentage of individuals classified to each region (rows) assigned to each inferred early-life region (columns). Percentages of residents (capture region matching the inferred early-life region) are highlighted in bold.

| Collection region | Reassigned region (%) |  |  |  |
| --- | --- | --- | --- | --- |
|  | Estuary | South | SBI | Undermined |
| Estuary | **15.7** | 24.9 | 38.0 | 21.4 |
| South | 19.5 | **31.0** | 26.1 | 23.4 |
| SBI | 5.4 | 13.2 | **70.4** | 11.1 |
| North | 9.7 | 24.7 | 47.4 | 18.2 |

Inter-annual comparisons indicated broadly consistent patterns between 2022 and 2024 (S4 Fig). These results suggest that the observed spatial patterns are robust.

### Drivers of movement and regional residency

Movement probability was influenced by both survey timing and inferred early-life region (Table 4). Capelin were less likely to be classified as migrants during the spawning period than during the post-spawning period. During spawning, 58.6% of individuals were captured in regions matching their inferred early-life region, whereas only 24.9% of post-spawning individuals were captured in locations matching their inferred early-life region (Fig 4). Moreover, individuals with SBI early-life signatures exhibited a lower probability of being classified as migrants than those from other regions. No significant effects of sex or maturity were detected. Overall, these results indicate that inferred early-life region and seasonal timing influence capelin movement patterns.

**Table 4.**
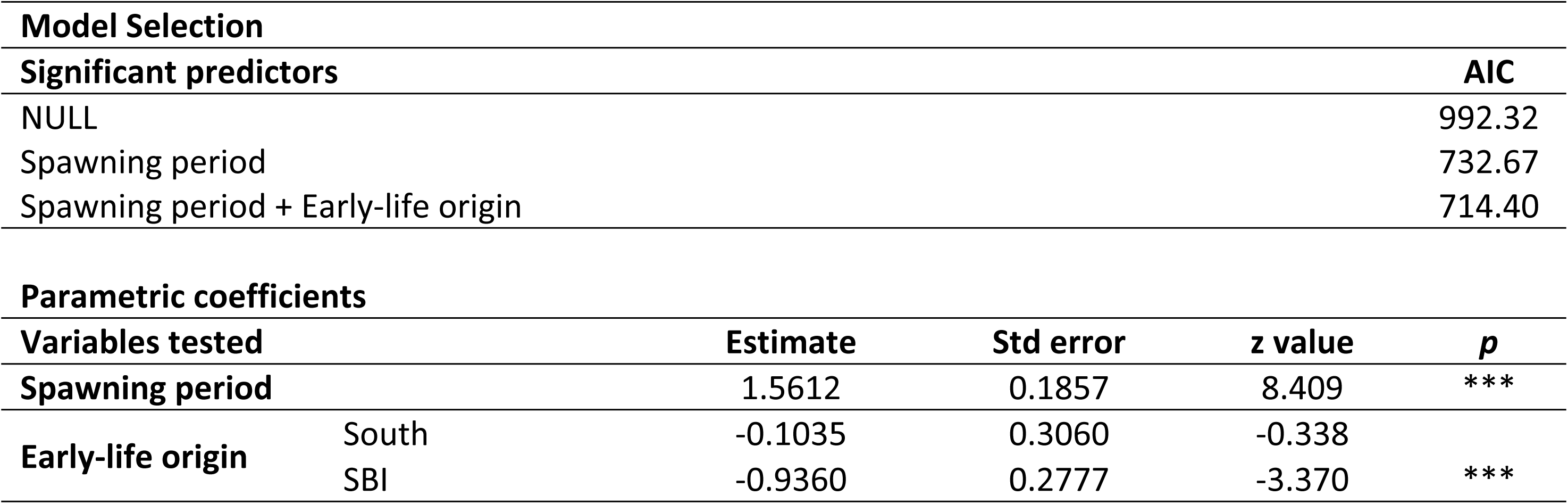
Generalised linear model (GLM) of the probability of capelin being classified as migrants, as a function of maturity, spawning period (during or after), sex, and inferred early-life region determined from otolith core analysis. Model selection was based on AIC. Coefficient estimates, standard errors, z-values, and p-values are presented. Significance levels are indicated as: *** p < 0.001, ** p < 0.01, * p < 0.05.

**Fig 4.**
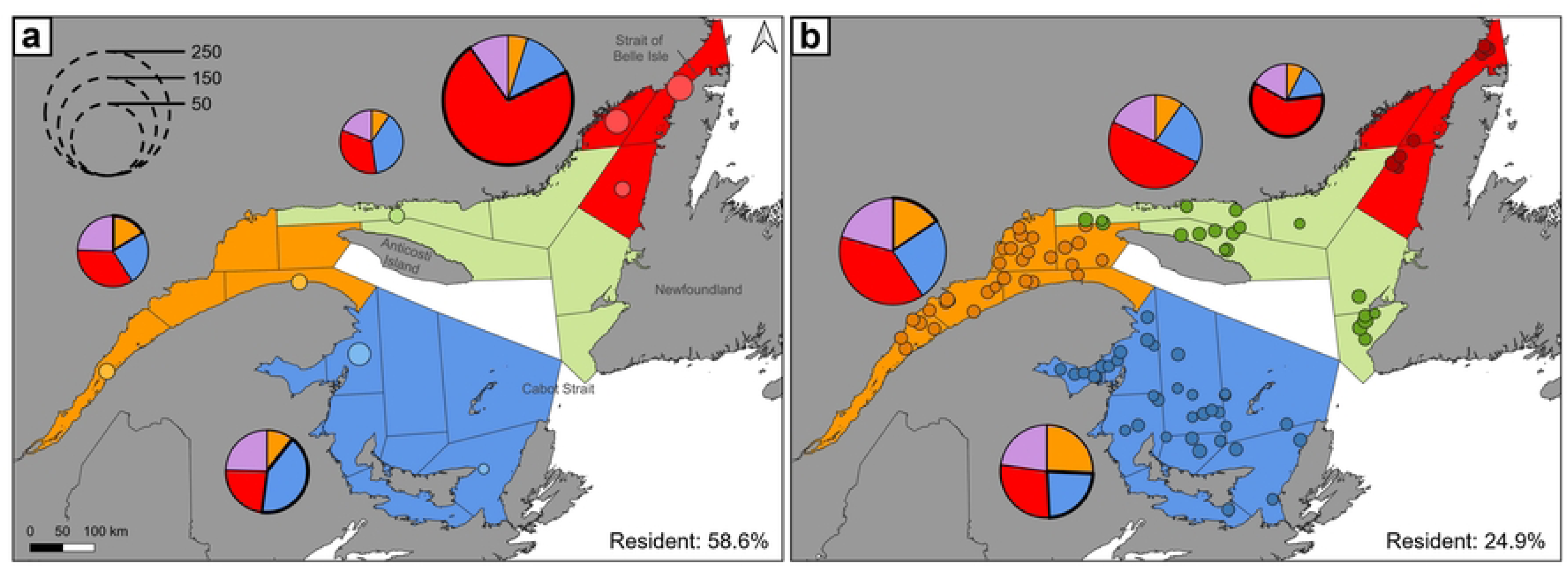
Spatial origin of capelin contingents sampled in the Gulf of St. Lawrence (a) during the spawning period and (b) after spawning. Colours indicate inferred early-life regions: orange for the Estuary, blue for the South, red for the Strait of Belle Isle and purple for an ‘undetermined’ region. Points represent capture locations and are proportional to the number of fish captured. In the pie charts, segments with thicker outlines represent resident individuals (those whose capture region matches their inferred early-life region), whereas those with thinner outlines indicate migrants. The size of each pie chart is proportional to the number of individuals captured in the area, as illustrated by the reference legend in the top left. Shapefiles used for map construction were obtained from Runfola et al. (2020). Map projections: EPSG:32198, NAD83 / Québec Lambert

## Discussion

The present study provides the first large-scale assessment of movement patterns of capelin in the GSL. By focusing on broad spatial structuring, results are robust to fine-scale spatial and temporal variability, which was deemed unlikely to bias the identified movement patterns. Throughout, the otolith core is treated as an integrated early-life regional signature, reflecting the late-larval to early-juvenile period rather than a precise natal origin, so that an individual’s ‘early-life region’ denotes where it resided during early life rather than where it was spawned. These patterns varied according to inferred early-life region and season. Individuals with SBI early-life signatures showed the strongest correspondence between early-life and capture regions, and this correspondence was stronger during spawning than post-spawning. More than half of the capelin captured in SBI carried a local early-life signature, whereas a higher influx of individuals with early-life signatures from elsewhere was observed in the Estuary and South regions. Overall, these results suggest that capelin connectivity in the GSL is characterised by a combination of widespread dispersal and partial regional residency. Such dynamics create a system in which some regions are well mixed while others seem predominantly self-sustained, potentially balancing demographic stability with gene flow.

### Patterns of capelin connectivity in the Gulf of St. Lawrence

Capelin in the GSL exhibit various patterns and levels of connectivity, and complex spatial dynamics. Overall, the population appears relatively well mixed, although mixing is less pronounced in the SBI than in other regions. At this level of connectivity, extensive gene flow limits genetic differentiation, consistent with earlier studies reporting broad genetic homogeneity across the region, albeit with some alleles occurring more frequently in specific areas (Cayuela *et al*., 2020; Dodson *et al*., 2007). During spawning, GSL capelin include both regionally resident individuals, representing more than half of those captured, and more dispersive individuals. This elevated correspondence between early-life and capture regions during the spawning season is consistent with regional spawning fidelity, which has previously been hypothesised for capelin (Kenchington *et al*., 2015), although the data from this study cannot establish a return to a precise natal site.

A notable proportion of individuals could not be confidently assigned to any of the three defined regions. This uncertainty may reflect individuals with early-life signatures from the northern zone, which lacks a sufficiently distinct elemental signature to allow high-confidence reassignments (Jac *et al*., 2026). Indeed, as this zone exhibits an intermediate chemical signature between the Estuary and the SBI, some northern individuals may have been assigned to one of these two regions based on the similarity of their otolith fingerprints. Alternatively, the presence of unassigned individuals may indicate contribution from additional, unsampled regions, outside the GSL, such as the Newfoundland or the Scotian shelves (Campana *et al*., 2000). Evidence for recurrent mixing between these stocks is provided by the occurrence of shared mitochondrial haplotypes across the GSL and the Newfoundland-Labrador Shelf (Cayuela *et al*., 2020; Dodson *et al*., 2007). Such exchanges are consistent with known oceanographic corridors. The SBI has been repeatedly reported as a migration route for capelin, facilitating exchanges between the GSL and the Newfoundland-Labrador Shelf (Cayuela *et al*., 2020; Kenchington *et al*., 2015), whereas the Cabot Strait serves as a corridor for pelagic fishes, including herring, mackerel, and cod (Brosset *et al*., 2019; Buren *et al*., 2014).

Connectivity patterns align with the dispersive behaviour capelin exhibit, including large-scale spawning migrations, seasonal offshore-inshore movements, and extensive feeding excursions (Carscadden *et al*., 1997; Crook *et al*., 2017; Jac *et al*., 2025; Rose, 2005). Such dynamics facilitate colonisation of new habitats, rescue effects, and metapopulation processes, while promoting gene flow and maintaining adaptive potential under changing environmental conditions (Cowen & Sponaugle, 2009; Hanski, 1998). The findings presented here extend beyond previous genetic studies and link observed elemental signatures with the ecological processes that sustain broad-scale connectivity in the GSL through the capelin’s life cycle.

### Spawning strategies: beach versus demersal

Capelin exhibit two main spawning strategies: beach spawning, during which adults roll onto the shoreline, and deeper demersal spawning (McQuinn *et al*., 2012). Beach spawning is better documented as it is easier to observe, and citizens can report beach-spawning events on GSL beaches (Boudreau *et al*., In press). In Newfoundland, these observations inform annual capelin stock assessments (DFO, 2025; Murphy, 2022; Murphy *et al*., 2021). However, focusing solely on beach-spawning events may overlook substantial demersal spawning (Bliss *et al*., 2023; Geoffroy *et al*., 2025), emphasising the need for complementary monitoring approaches capable of detecting both spawning modes. The relative reliance on these strategies varies with local thermal conditions (Davoren, 2013; Tripp *et al*., 2025), with demersal spawning contributing less when spawning is delayed (Tripp *et al*., 2025). Previous research showed that core otolith chemistry may differ between these spawning modes (Tripp & Davoren, 2025). In the present study, spawning sites were not directly sampled, and cores from beach and demersal spawners were not compared. Distinguishing the contributions of individual sites or spawning modes within a region is therefore beyond the resolution of the broad-scale, edge-based framework used here. For the same reasons discussed above, which prevent to infer finer-scale inter-annual and among-site variation from the present analysis, our inferences are confined to the among-region scale, at which spatial structuring is pronounced (Jac et al., 2026).

During the spawning season, temperatures in the SBI, the region characterised by the coldest surface waters in the GSL, remain within the preferred range for capelin (Boudreau *et al*., In press). Bottom temperatures between −1 and 1 °C and the presence of a thick CIL, typically between 25 and 75 m, appear to provide extensive habitat suitable for both beach and demersal spawning (Galbraith *et al*., 2025). Capelin captured in this region are, on average, larger and older than those from other regions (S5 Fig), consistent with patterns reported by Lehoux *et al*., (2022). These results indicate a strong local contribution (i.e., substantially higher than in other regions; Table 3) to spawning by resident fish and/or mature fish returning to the SBI. This regional residency could reflect fitness advantages, including reducing habitat search costs, lowering predation risk, and increasing mate encounter rates (Swearer *et al*., 2002).

Elsewhere in the GSL, surface waters are warmer, and the CIL is thinner (Boudreau *et al*., In press; Galbraith *et al*., 2025). In the southern GSL, bottom temperatures remain relatively cold, generally between 0 and 2 °C except in the Northumberland Strait (Galbraith *et al*., 2025). These low temperatures, combined with the presence of shallow shelf areas, mostly shallower than 75 m, appear to provide extensive habitat suitable for demersal spawning. Moreover, given the strong seasonal variability in temperature (Galbraith *et al*., 2025), capelin may favour earlier spawning at cooler subtidal sites, allowing embryos to hatch earlier and larvae to benefit from an extended growth period prior to overwintering, ultimately enhancing survival (Tripp *et al*., 2025). Although beach-spawning events are rarely reported in these areas (Boudreau *et al*., In press; Jac *et al*., 2025), this likely reflects limited observation rather than true absence. Supporting this, systematic surveys, including mackerel egg sampling, document larval capelin in Chaleur Bay and St. George’s Bay (DFO, *unpublished data*), and length-frequency data indicate abundant small capelin consistent with local nursery production (Lehoux *et al*., 2022). These findings suggest that the southern GSL provides important spawning and nursery habitats, generating a strong early-life signal despite the scarcity of reported beach-spawning events.

In the Estuary, where most previous studies within the GSL have been conducted (e.g., Bailey *et al*., 1977b; Jacquaz *et al*., 1977; Lazartigues *et al*., 2016; Marchand *et al*., 1999), a major nursery area has been identified for capelin (Ouellet *et al*., 2013). The Estuary is relatively warm during the spawning season (Galbraith *et al*., 2025), which limits the potential for demersal spawning while still allowing beach spawning to occur (Davoren, 2013; Boudreau *et al*., In press; Jac *et al*., 2025). Although the Estuary is of high ecological importance for capelin during the late-juvenile stage, results indicate that other regions of the GSL also play essential roles at earlier life stages.

### Larval drift and juvenile/adult migrations

Connectivity in GSL capelin may involve both larval drift and the seasonal migrations of juveniles and adults. During their relatively short larval phase, typically 1.5-3 months, depending on temperature (Bailey *et al*., 1977b; Jacquaz *et al*., 1977), capelin larvae largely occupy the upper water column, where their distribution is strongly influenced by prevailing currents (Dalley *et al*., 2002; Taggart & Leggett, 1987). Retention has been documented in localised areas such as the Estuary (Ouellet *et al*., 2013) and coastal bays (Carscadden, 1979; Grégoire & Girard, 2014; Sirois *et al*., 2009), indicating that circulation can concentrate larvae regionally. However, strong currents can transport larvae from the Estuary into the southern GSL (El-Sabh, 1976; Galbraith *et al*., 2025). The Labrador Current, entering through the SBI, also likely facilitates dispersal across the broader GSL system (Shaw & Galbraith, 2023). Despite these oceanic processes, the short larval duration relative to the spatial scale of the GSL makes larval drift unlikely to be the main driver of large-scale connectivity, although it might contribute to local recruitment variability and retention.

Beyond larval dispersal, juvenile and adult capelin undertake pronounced seasonal migrations, reflected in varying proportions of migrants and residents between spawning and post-spawning periods. The Estuary appears to receive substantial inflow, as many individuals sampled there carry early-life signatures from elsewhere in the GSL. Such movements require active migration and are consistent with patterns observed in the Saguenay Fjord, where individuals migrated up to 300 km during their first year (Lazartigues *et al*., 2016). Through these migrations, capelin can track favourable conditions in terms of prey availability, thermal environments, and predation pressure (e.g., Fall *et al*., 2018; Ingvaldsen & Gjøsæter, 2013; Rose, 2005). From offshore wintering areas, capelin migrate inshore to spawn around July, covering distances of up to 350 km (Carscadden *et al*. 2013; Templeman, 1948). Males typically arrive first and remain throughout the spawning period, whereas females release eggs and then return to deeper waters (Davoren, 2013). Foraging migrations are closely linked to prey distribution, including copepods, amphipods, and euphausiids (Aarflot *et al*., 2020; Dalpadado & Mowbray, 2013), and are also influenced by environmental factors such as sea temperature and ice cover (Mowbray, 2002; Ogloff *et al*., 2020). In the SBI, the mixing of the Labrador Current and GSL outflow creates favourable trophic and thermal conditions, enhancing prey availability and extending feeding seasons (Cyr & Larouche, 2015; Laliberté & Larouche, 2023), which may explain the higher proportion of resident individuals in this region. Overwintering strategies are similarly shaped by circulation and thermal regimes. In the GSL, the CIL and the strong seasonal variability of the southern GSL restrict habitat suitability (Chamberland *et al*., 2022; Galbraith *et al*., 2025), guiding seasonal movements toward deeper thermal refuges in the Cabot Strait, the Estuary, and the SBI, consistent with overwintering patterns observed off the northeast coast of Newfoundland (Carscadden, 1984; Minet & Perodou, 1978; Winters & Carscadden, 1978).

### Integrating connectivity and behaviour into capelin management

While capelin stocks in areas such as the Barents Sea, the Newfoundland-Labrador Shelf, and Iceland are managed with explicit recognition of their ecological role (Singh *et al*., 2025a, 2025b), such integration through the Ecosystem Approach to Fisheries (e.g., Garcia & Cochrane, 2005; Koen-Alonso *et al*., 2019) is currently lacking in the GSL. The findings of this study have multiple implications for management, including providing context for ecosystem-based considerations, clarifying the risk of localised depletion, improving interpretation of fishery and survey indicators, and guiding monitoring and survey design.

The coexistence of dispersive and regionally resident strategies generates a portfolio effect, distributing reproductive effort across space and time and buffering populations against environmental variability, predator outbreaks, and localised reproductive failures (Schindler *et al*., 2010, 2015). Similar patterns of behavioural diversity have been observed in mackerel in Canadian waters (Piczak *et al*., *Under review*). Maintaining this behavioural diversity is therefore essential for long-term population stability, as its erosion has been linked to collapses in small pelagic fishes (Price *et al*., 2021). Reliable estimates of early-life regional composition and population structure further improve understanding of how behavioural and spatial diversity supports the persistence of this heavily exploited stock under environmental stressors, such as altered flows or episodic disturbances (Sturrock *et al*., 2020).

Although these patterns do not define distinct assessment units, they have clear management implications. Spatial heterogeneity can increase the risk of localised depletion, bias survey-based indices, and complicate interpretation of temporal trends (Randon *et al*., 2021). The SBI was identified as a region with high local production and substantial commercial fishing effort, making it particularly susceptible to localised depletion and highlighting the need for spatially explicit management measures. Heterogeneous connectivity also affects the interpretation of fishery and survey indicators: temporal changes in catch-per-unit-effort (CPUE) on the west coast of Newfoundland (Boudreau *et al*., In press) may partly reflect shifts in connectivity rather than overall stock productivity, and bottom-trawl surveys, covering only part of the GSL, cannot be interpreted independently without accounting for regional movements and mixing. Incorporating connectivity information therefore reduces the risk of biased stock assessments due to localised signals (Le Pape et al., 2020).

Survey design and monitoring programmes should aim for broad spatial coverage to capture the full spectrum of population composition, life stages, and regional contributions, ensuring that derived indicators reflect stock-wide dynamics rather than localised subunits. In this context, the present study provides empirical evidence of substantial connectivity across the GSL, a feature previously suspected but undocumented. By linking individual movement histories and inferred early-life regions to spatial population structure, these findings enhance understanding of capelin stock dynamics, inform ecosystem-based management, and support the development of more robust survey designs and future research directions.

### Uncertainty and limitations

Otolith elemental composition reflects a complex interplay between environmental conditions (e.g., temperature, salinity, productivity) and intrinsic physiological processes (e.g., growth rate, metabolism), and both must be accounted for when reconstructing environmental histories (Hüssy et al., 2020; Miller & Hurst, 2020; Barnes & Gillanders, 2013; Reis-Santos et al., 2012). A central question for the present study is which life-history window the otolith core records. In capelin, larval otoliths display markedly elevated trace-element concentrations in both the pre-hatch and post-hatch regions, corresponding to the first months of life (Coussau et al., 2026; Davoren & Halden, 2014; Fink-Jensen et al., 2022; Tripp et al., 2020), yet these elevated concentrations were not recovered in the adult cores analysed here. The most parsimonious interpretation is therefore that the core does not capture the larval period as strictly defined, but rather an integrated window spanning the late-larval to early-juvenile (settlement) phase, the underlying mechanisms remaining unresolved (Coussau et al., 2026). Importantly, this uncertainty about the precise period recorded does not diminish the value of the core signal: even without identifying where individuals were spawned, the core integrates a formative early-life window, encompassing the growth and survival processes that shape recruitment and future stock dynamics, and therefore remains ecologically meaningful as an integrated early-life regional signature rather than a strict natal marker.

A further consideration is temporal stability. Multi-year stability of region-specific signatures has been demonstrated on the otolith edge, which reflects the adult capture environment (Gauthier et al., 2024; Jac et al., 2026), but the stability of core signatures within regions was not directly tested here. This distinction matters because the early-life window is laid down by larvae and early juveniles, whose physiology and habitat differ from those of the adults on which the regional baseline was validated, so edge stability cannot be assumed to carry over directly to the core. At finer scales, inter-annual variation in larval otolith chemistry and substantial among-site heterogeneity have both been documented for capelin and can reduce discrimination success (Coussau et al., 2026; Tripp et al., 2020). Together, these constraints confine our inferences to the broad, region-level scale and preclude individual-level natal assignment.

Despite these limitations, several lines of evidence indicate that the broad-scale patterns reported here are robust. First, the assignments are anchored to otolith-edge signatures whose multi-year stability and strong spatial contrast are established, with among-region differences consistently exceeding inter-annual variation, and which Jac et al. (2026) attributed primarily to large-scale, temporally persistent drivers rather than to ephemeral local conditions; because such drivers are stable over time, the among-region contrast they generate was likely expressed during the early-life period as well, lending indirect support to the use of the core signal for broad regional assignment. Second, the high concordance between age-stratified and pooled models (80.5% for directly aged fish; 97.0% for the age-supplemented dataset) indicates that cohort effects have limited influence on assignments, reflecting internal consistency across cohorts. Third, the consistently low proportion of ‘undetermined’ individuals is informative: because the three regions are strongly chemically differentiated (Jac et al., 2026), a core signature incompatible with all three falls below the assignment threshold and is classified as ‘undetermined’ rather than forced into an incorrect region, so the low frequency of such individuals indicates that most core signatures are compatible with a single regional signature.

Finally, the applicability of this framework is system-dependent. The environmental stability of the GSL during summer relative to the species’ short lifespan, together with the broad spatial scale of the analysis relative to the scale of physiological and environmental variability, supports the use of edge-to-core reassignment to detect large-scale connectivity. Caution is warranted, however, when extending the approach to finer spatial scales, to more dynamic environments, or to longer-lived species.

## Acknowledgments

We extend our sincere gratitude to everyone involved in capelin sampling and otolith collection at Fisheries and Oceans Canada. We are especially thankful to Laurence Lévesque, Mélanie Boudreau, Catherine Fortin-Tanguay, Chantal Méthot, and Rachel Mailhot for their work on otolith extraction and preparation. We also greatly appreciate the support of Félix Gagnon and Dany Savard (LabMaTer, UQAC) for their assistance with the LA-ICP-MS analyses, as well as that of Anne-Lise Fortin (UQAC) for overall project logistics. We dedicate this work to the late Stéphane Plourde, a collaborator and friend who played a key role in conceptualising this study.

## Ethics statement

All otoliths analysed in this study were obtained from fish collected during routine DFO scientific surveys or from commercial fishing operations conducted in accordance with applicable national and institutional regulations. No live animals were handled or euthanised specifically for this study, and all samples were collected as part of ongoing monitoring or harvesting activities. Consequently, no additional animal care or ethics approval was required.

## Data availability statement

Data files used for this study are available upon request from the Université du Québec à Rimouski collection in the Borealis data repository (https://doi.org/10.5683/SP4/X0CVNA). Shapefiles can be found in www.geoboundaries.org and https://www.naturalearthdata.com/.

## Author contributions

R.J., E.V.B, M.B, D.R., P.B., O.L.P., and P.S. conceptualised the study. Data were generated by R.J., P.S, E.V.B. and M.B. R.J. conducted data analysis and led the project under the supervision of D.R., P.B., O.L.P., and P.S. R.J. drafted the manuscript, with all authors contributing to result discussions, manuscript revisions, and approving the final version.

